# Optogenetic inhibition and activation of Rac and Rap1 using a modified iLID system

**DOI:** 10.1101/2020.12.11.421990

**Authors:** R. Nick Elston, Michael Pablo, Frederico M. Pimenta, Klaus M. Hahn, Takashi Watanabe

## Abstract

The small GTPases Rac1 and Rap1 can fulfill multiple cellular functions because their activation kinetics and localization are precisely controlled. To probe the role of their spatio-temporal dynamics, we generated optogenetic tools that activate or inhibit endogenous Rac and Rap1 in living cells. An improved version of the light-induced dimerization (iLID) system was used to control plasma membrane localization of protein domains that specifically activate or inactivate Rap1 and Rac (Tiam1 and Chimerin for Rac, RasGRP2 and Rap1GAP for Rap1). Irradiation yielded a 50-230% increase in the concentration of these domains at the membrane, leading to effects on cell morphodynamics consistent with the known roles of Rac1 and Rap1.

## Introduction

The small GTPases Rap1 and Rac act as molecular switches in multiple cellular signaling circuits [1]. They interact with downstream effectors when they are in a GTP-bound ‘on’ state, and hydrolyze the bound GTP to turn themselves off. Rap1 is a Ras family small GTPase that is important in cell growth, differentiation, cell adhesion and the MAP-kinase cascade [2,3]. Rac1 is a Rho family GTPase that regulates diverse cell functions, including cell growth, motility, cytoskeletal reorganization and apoptosis [1]. The activity of both GTPases is controlled by GTPase activating proteins (GAPs) that inactivate the proteins by accelerating GTP hydrolysis, and guanine nucleotide exchange factors (GEFs) that produce the activated conformation by catalyzing GTP binding [4].

In this paper we describe a method to optogenetically control fragments of GEFs and GAPs to activate or inhibit endogenous Rac and Rap1. To our knowledge, Rap1 activity and Rac GAP activity have not previously been controlled optogenetically. Published strategies to control Rac include incorporation of the light-responsive LOV domain from *Avena Sativa* in Rac1 to allosterically or sterically control the active site [5,6], and driving GEFs to the plasma membrane through light-induced heterodimerization (using the dimerizers CRY/CIBN [7], PhyB/PIF [8], LOV2 fragment/ePDZ [9], and LOV2/stringent starvation protein (ssr) A/B [10]). Here we used improved LOV-induced dimerization [10], a published, light-induced dimerization approach, to control membrane localization of the Rap1-specific GEF RasGRP2 and GAP Rap1GAP. Rac activity was controlled using a fragment of the GAP Chimerin [1].

The iLID approach takes advantage of a high affinity interaction between the bacterial peptide SsrA and the bacterial protein SspB [10]. LOV2 is modified to incorporate SsrA in the carboxyl terminal Jα helix, where it is exposed only upon irradiation. A localization sequence is used to attach this SsrA-LOV fusion to the plasma membrane [10]. When blue light irradiation exposes the SsrA, proteins fused to SspB bind to it, and therefore accumulate at the membrane.

## Materials and methods

### Plasmid construction

pLL7.0: Venus-iLID-CAAX (from KRas4B) and pLL7.0: mTiam1(64-437)-tgRFP-SspB wild type were a gift from Brian Kuhlman (Addgene plasmid # 60411; http://n2t.net/addgene:60411; RRID:Addgene_60411, and Addgene plasmid # 60417; http://n2t.net/addgene:60417; RRID:Addgene_60417). To optogenetically control Rac1 and Rap1 activity, the following specific GEF and GAP constructs were used: Rac1 GEF, mouse Tiam1 (1033-1406 amino acid in NP_001139358.1); Rac GAP, human β-Chimerin (141-332 amino acid in NP_001035025.1); Rap1 GEF, mouse RasGRP2 (2-424 amino acid in AAC79697.1), Rap1 GAP (75-415 amino acid in AAH54490.1). These catalytic domains were subcloned into a pmVenus-C1 vector together with a nuclear export signal (NES) fused to SspB wild type, a tandem ribosomal skip sequence, flag tag, and iLID CAAX by Gibson assembly (New England Biolabs, MA, USA). The tandem skip sequence was used to express the two components of the iLID system from a single mRNA, as previously described [11]. To generate the control plasmid, pmVenus-mTiam1-SspB-T2AP2A-iLID-CAAX was digested with HindIII and self-ligated to excise the entire Dbl homology domain and half of the Pleckstrin homology domain. The 16 amino acids of the Lyn kinase amino terminus (MGCIKSKRKDNLNDDE), which undergo lipid modification in cells, were fused to the N terminus of mCherry (nLYN-mCherry) and used as a membrane-localized fluorescent marker to monitor the plasma membrane.

### Cell culture and transfection

HeLa cells were obtained from American Type Culture Collection (ATCC CCL-2). HeLa and LinXe cells were maintained in DMEM containing 10% fetal bovine serum (Gemini Bio-Product, CA, USA) and 2 mM GlutaMax (Thermofisher Scientific, MA, USA). Fugene 6 transfection reagent (Promega, WI, USA) was used for transient transfection according to the manufacturer’s instructions. In brief, 200 ng of nLYN-mCherry and 500 ng of iLID plasmid were mixed with 2.1 ul (1 to 3 ratio) of Fugene 6 in 125 ul of Opti-MEM medium (Thermofisher Scientific). After 10 minutes incubation at room temperature, the mixture was overlaid onto cells cultured on a fibronectin-coated coverslip (Sigma Aldrich) in a 6-well plate. The cells were imaged within 20 hours after the transfection to avoid overexpression. Identical culture and transfection methods were used in local activation experiments, except the concentration of iLID plasmid DNA was doubled for the Chimerin regulator.

### Immunoblot analysis

Transfected LinXe cells were washed with PBS twice and dissolved in SDS-PAGE sample buffer. Total proteins were separated in SDS-PAGE and transferred to a PVDF membrane using Transblot turbo (Bio-Rad Laboratories, CA, USA). After blocking the membrane with 0.3% skim milk in TBS-T, the membrane was incubated with primary antibody for either 1 hour at room temperature or overnight at 4°C, and then with secondary antibody for 2 hours at room temperature. The membrane was washed with TBS-T three times (5 minutes each) after the incubation with the antibodies. The following antibodies were used: Anti-GFP antibody (JL-8 from Takara, Shiga, Japan), anti-Flag antibody (M2 from Sigma-Aldrich, MO, USA), and anti-mouse IgG DyLight conjugate (Cell Signaling Technology, MA, USA). The fluorescent images of the membrane were captured using the ChemiDoc MP imaging system (Bio-Rad Laboratories).

### Imaging

Before imaging, each coverslip was transferred to a home-made chamber and the medium changed to imaging medium (Phenol red-free Ham’s F12 (Caisson Labs, UT, USA) supplemented with 2% fetal bovine serum and 15 mM HEPES (Thermofisher Scientific)). The imaging system and scope for local activation were as described previously [12]. Cells were exposed to blue light by irradiating a circle of area 1089-1175 µm^2^ using 0.810 µW intensity (as measured by placing a Thorlabs S121B power meter head at the microscope revolver, before the objective) with six seconds of exposure per cycle. This resulted in 4.9 µJ of energy per irradiation cycle.

### Image processing and quantification

Images were threshholded, corrected for photobleaching [13], and cell masks were generated using the software MovThresh [14]. The region of the edge proximal to irradiation (ROI) was defined as follows: A binarized image of the irradiation spot was created. A dilation of the binarized image was calculated using a disk structural element with a 40 pixel radius and 8-line-segment resolution (MATLAB function strel(‘disk’, 40, 8)) (MATLAB 2017b, Natick, MA). Separately, a binarized cell mask was created and the cell boundary was obtained. A dilation of the binarized cell boundary was calculated using a disk structural element with a 4 pixel radius (MATLAB function strel(‘disk’, 4). Pixels of the dilated boundary outside of the binarized cell mask were thrown out. The region of the edge proximal to irradiation was taken as the overlapping pixels between the dilation of the binarized irradiation spot, and the dilated binarized cell mask. The distal regions were the remaining pixels of the dilated binarized cell mask, that did not overlap the dilated binarized irradiation spot. For the two regions, the mean mVenus intensity was measured over time and normalized to three-frame pre-stimulation means. For protrusion/retraction analysis, a linescan was drawn through the cell starting from the irradiation zone, out through the other side. The movement of the cell edge along the linescan was normalized by subtracting the three-frame pre-stimulation mean.

## Results

Light-induced dimerization enables protein recruitment to the plasma membrane with precise kinetics and subcellular localization. The successful use of a Rac GEF fragment to *activate* Rac in previous work (Tiam1 DH/PH domain [10]) inspired us to explore Rac and Rap1 *inactivation* using GAPs - specifically Chimerin [15] for Rac, and Rap1GAP for Rap1 [16,17]. We also sought to activate Rap1, so applied iLID to a domain from the GEF RasGRP2. Each regulatory protein fragment was selected because it was specific for our targeted protein [15,17–19], and the previously described Tiam1 fragment was used as a control. Fragments of the GAPs and GEFs were fused to both the fluorescent protein mVenus and to SspB (**Figs 1a and 1b**). These fusions were expressed together with flag-tagged LOV2-SsrA using a modified viral ribosomal skip sequence to express separate genes from one construct, producing a more consistent expression ratio for the two proteins. Immunoblot analysis of cell lysates from LinXe cells confirmed that the two chains were expressed separately, and that there was relatively small variation in the expression ratio of the two chains (**Figs 1c and 1d**).

**Fig 1.**
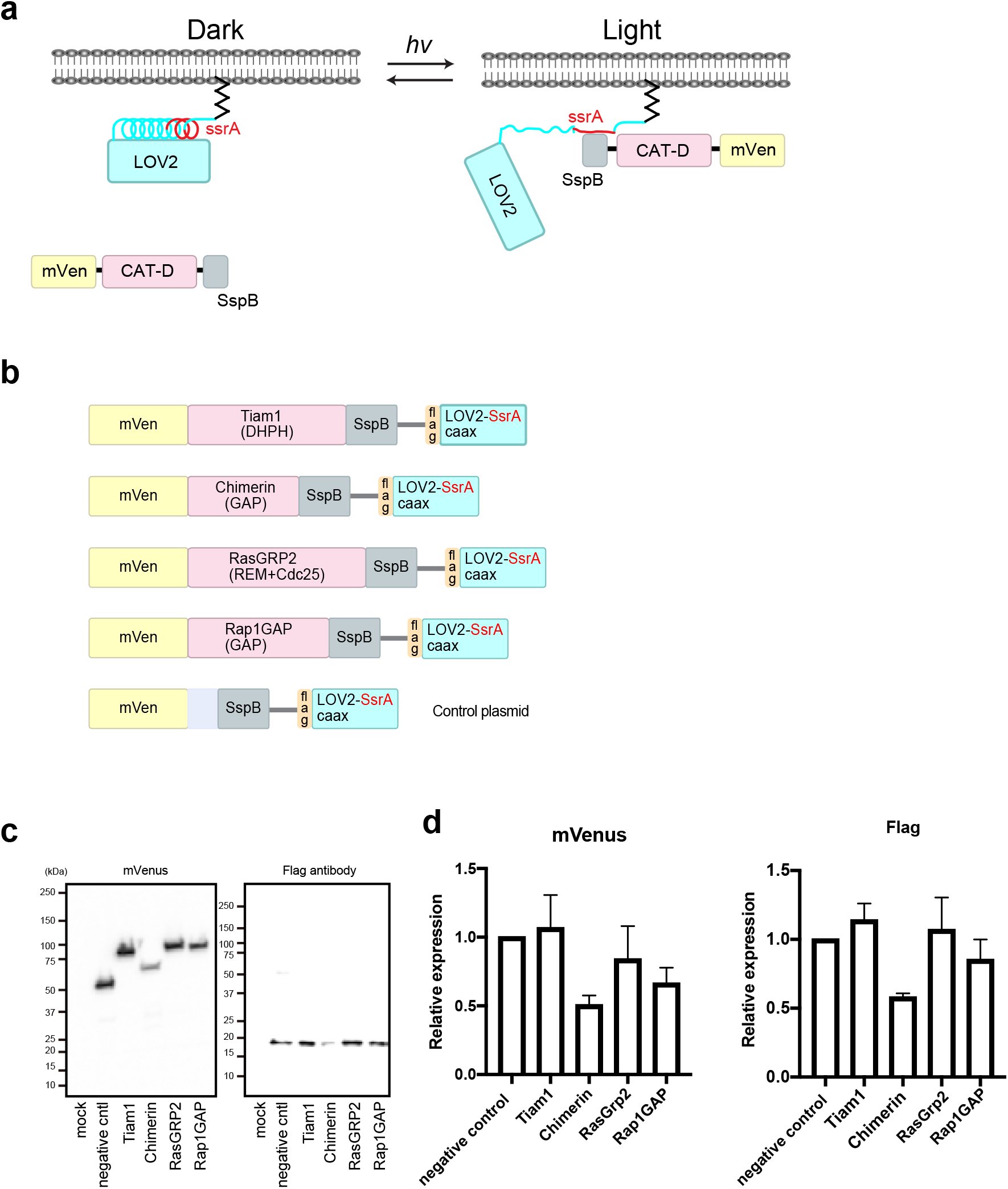
Engineered iLID constructs for optogenetic control of Rap1 and Rac activity. (a) Proteins for reversible photoinduced recruitment of GEF or GAP catalytic domains (CAT-D) to the plasma membrane. (mVen = mVenus fluorescent protein) (b) Design of Rac1 and Rap1 iLID-based regulators. The catalytic domains of Tiam1 (Rac activator), Chimerin (Rac inactivator), RasGRP2 (Rap1 activator), and Rap1GAP (Rap1 inactivator) were inserted as shown. A ribosomal skip sequence was used to produce separate expression of the two components from one gene. (c) Immunoblots showing separate expression of both components. mVenus was visualized using an anti-GFP antibody. (d) Densitometry of the gels in panel c, showing relatively fixed ratio of the two components.

To verify light-induced translocation of the catalytic domains to the plasma membrane we irradiated a circular spot at the edge of HeLa cells, alternating six seconds of irradiation with six seconds of darkness for a total of 5 minutes. During this process images were obtained every 10 seconds, beginning 5 minutes prior to irradiation. The mean mVenus intensity was measured in a band extending 1.3 microns inward from the cell edge. We compared the average intensity of the edge within 17.5 microns of the spot center to the remainder of the edge (**Figs 2a, 2b**). The intensity of mVenus in the proximal region increased 50-230% during irradiation, whereas the intensity of the region distal to irradiation was minimally affected (**Fig 2b)**. Kymograph analysis of cells expressing Rap1GAP showed that light-induced retraction correlated with increased mVenus intensity at their membranes (**Fig 2d**). Edges proximal to the irradiation spot retracted after irradiation, while regions distal to irradiation were not affected (**Fig 2e**).

**Fig 2.**
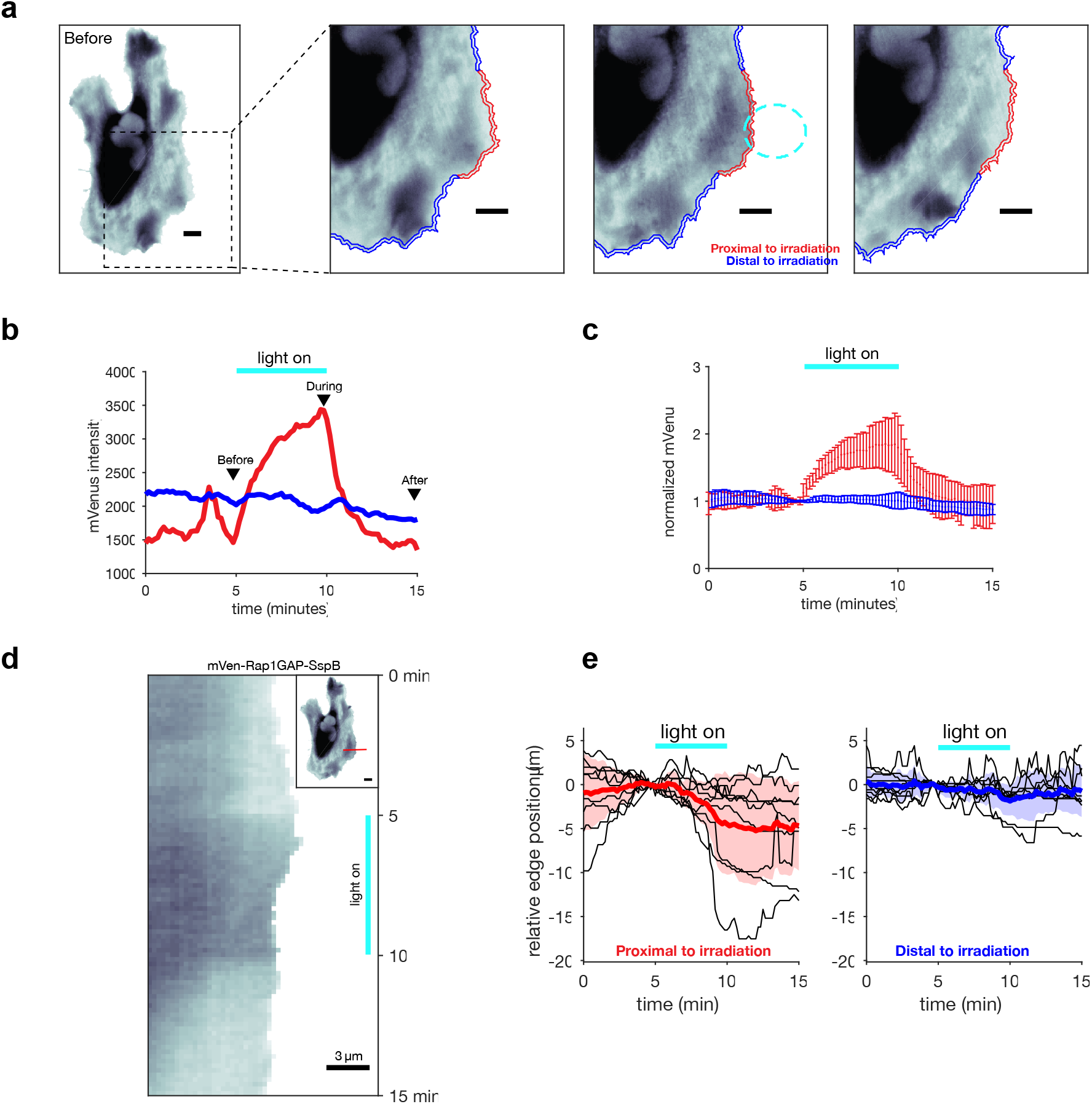
Irradiation of mVenus-Rap1GAP-SspB. (a) Example HeLa cell before, during, and after localized irradiation. Cell edge portions proximal to irradiation (a17.5 microns from irradiation center) are shown in red. All other portions of the cell were considered distal to irradiation (blue), extending all the way around the cell. Scale bars, 10 µm. (b) The mVenus intensity from panel a averaged within regions proximal (red) and distal (blue) to the irradiation zone. (c) Membrane translocation responses normalized to pre-stimulation, mean±1s.d. (n=9 cells). (d) mVenus kymograph of the HeLa cell shown in panel a, proximal to irradiation, illustrating Rap1GAP can induce retraction even of a protruding cell region, as shown here. (e) Protrusion quantified for Rap1GAP, normalized to pre-stimulation. Individual cells shown as solid black lines, mean±1s.d. shaded in color (n=9 cells).

We next characterized light-induced protrusion or retraction for all the constructs (**Fig 3, supplemental movies S1-S4**). Each regulator produced a statistically significant response near the site of irradiation (solid red line), while distal regions exhibited constitutive protrusion-retraction cycles unchanged by irradiation, with an average of zero edge displacement (blue solid line). The negative control (mVenus-SspB) exhibited no protrusion/retraction response despite strong recruitment. As expected, the GEFs Tiam1 and RasGRP2 produced protrusion, while the GAPs Chimerin and Rap1GAP caused retraction. The regulators also produced more complex changes in either ruffling or the frequency of protrusion/retraction cycles.

**Fig 3.**
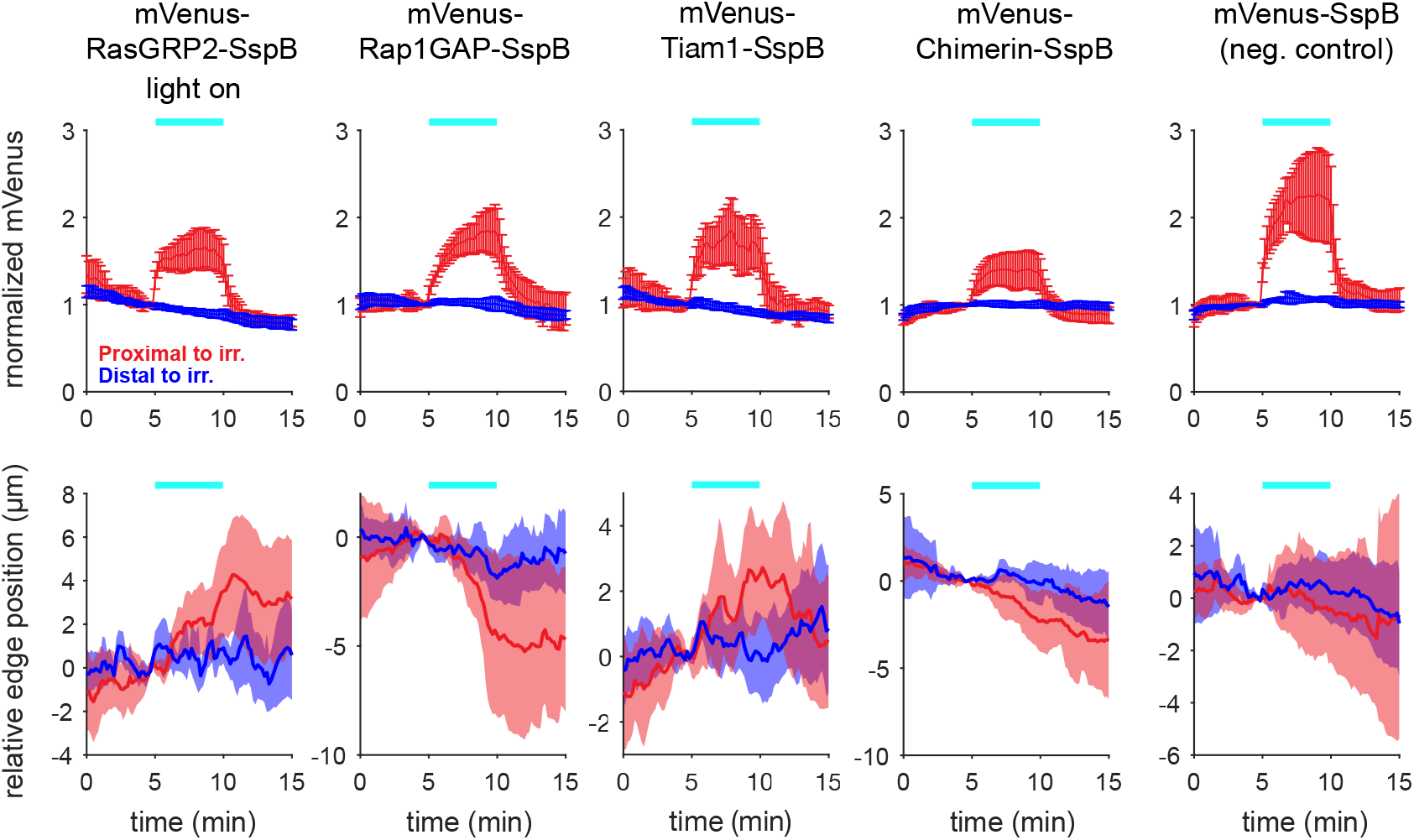
Light-dependent translocation (top) of GEF and GAP domains, with resultant effects on protrusion/retraction (bottom). The known Tiam1 construct and m-Venus SspB are included as controls. All curves show a 95% confidence interval; RasGRP2, n=10; Rap1GAP, n=9; Tiam1, n=5; Chimerin, n=8; negative control, n=9.

In addition to membrane protrusion, recruitment of Tiam1 promoted the formation of vesicles near the area of irradiation (**Supplemental S1 Fig**). This result is consistent with previous findings that activation of Rac1 induces endocytosis [20], providing additional evidence that Tiam1 manipulates Rac1 activity.

## Discussion

Optogenetic tools are powerful for studying biological processes because they afford precise manipulation of signaling in both time and space. We report here optogenetic activation and inactivation of Rap1, as well as inactivation of Rac. The single-gene tools presented here can act rapidly and reversibly. The unwinding of the LOV alpha helix with exposure of the dimerization domain occurs in less than a second [21–23]. Activation is then limited by diffusion of the activated regulator to the nearby membrane. We saw movement of the regulatory construct to the membrane within 10 seconds, and a protrusion or retraction response in less than a minute. Consistent with its reported roles [24], Rap1 activation induced membrane protrusion and inactivation produced retraction. Also consistent with its known roles, inhibition of Rac1 produced retraction. This data shows for the first time that photomanipulation of Rap1 on a short time scale is sufficient to cause quantifiable changes in cell membrane dynamics.

Interestingly, while recruitment of RasGRP2 and Chimerin caused membrane changes that persisted after turning off the blue light, Tiam1 effects were more transient, with protrusion returning to levels like those at distal regions soon after irradiation ceased (**Supplemental movies**). This may be due to negative feedback pathways for Rac [25–27].

The extent of response to irradiation was heterogeneous, with some cells responding substantially more or less than the average. This could be due to heterogeneous expression of the regulators and/or because the cells were polarized. In polarized cells, signaling pathways or cytoskeletal configurations can favor protrusion or retraction at a given region of the cell edge. Despite this heterogeneity, we observed statistically significant translocation and protrusion/retraction phenotypes for each regulator.

We hope these tools will provide a valuable means to control Rap1 and Rac in the wide range of cell behaviors that they regulate.

## Acknowledgements

We thank Wolfgang Bergmeier (University of North Carolina at Chapel Hill), Brian Kuhlman (University of North Carolina at Chapel Hill) and Kozo Kaibuchi (Nagoya University) for sharing cDNA and helpful suggestions.

## Supporting Information

**S1 Fig. Light-induced vesicle formation in a cell expressing the Tiam1 construct**. Blue ellipse indicates location of blue light irradiation. Red arrow indicates vesicles.

**S1 Movie. Chimerin**. The white dot indicates when the cell was irradiated. Note reversible light-induced retraction and induction of cell surface ruffles. Movie covers 60m real time compressed into 17 seconds, with an interval between frames of 30s real time.

**S2 Movie. Rap1 GAP**. The white dot indicates when the cell was irradiated. Note reversible light-induced retraction and increase in the frequency of cell edge protrusion/retraction cycles. Movie covers 60m real time compressed into 17 seconds, with an interval between frames of 30s real time.

**S3 Movie. RasGRP2**. The white dot indicates when the cell was irradiated. Note reversible, light-induced retraction and increase in the frequency of protrusion/retraction cycles. Movie covers 60m real time compressed into 17 seconds, with an interval between frames of 30s real time.

**S4 Movie. Tiam1 control**. The white dot indicates when the cell was irradiated. Note reversible induction of ruffling and protrusion. Movie covers 60m real time compressed into 17 seconds, with an interval between frames of 30s real time.

## References

1. Ridley AJ. Rho GTPases and actin dynamics in membrane protrusions and vesicle trafficking. Trends Cell Biol. 2006;16: 522–529. doi:10.1016/j.tcb.2006.08.006

2. Kooistra MRH, Dubé N, Bos JL. Rap1: a key regulator in cell-cell junction formation. J Cell Sci. 2007;120: 17–22. doi:10.1242/jcs.03306

3. Dillon TJ, Carey KD, Wetzel SA, Parker DC, Stork PJS. Regulation of the small GTPase Rap1 and extracellular signal-regulated kinases by the costimulatory molecule CTLA-4. Mol Cell Biol. 2005;25: 4117–4128. doi:10.1128/MCB.25.10.4117-4128.2005

4. Reiner DJ, Lundquist EA. Small GTPases. WormBook. 2018;2018: 1–65. doi:10.1895/wormbook.1.67.2

5. Dagliyan O, Tarnawski M, Chu P-H, Shirvanyants D, Schlichting I, Dokholyan NV, et al. Engineering extrinsic disorder to control protein activity in living cells. Science. 2016;354: 1441–1444. doi:10.1126/science.aah3404

6. Wu YI, Frey D, Lungu OI, Jaehrig A, Schlichting I, Kuhlman B, et al. A genetically encoded photoactivatable Rac controls the motility of living cells. Nature. 2009;461: 104–108. doi:10.1038/nature08241

7. Valon L, Marín-Llauradó A, Wyatt T, Charras G, Trepat X. Optogenetic control of cellular forces and mechanotransduction. Nat Commun. 2017;8: 14396. doi:10.1038/ncomms14396

8. Levskaya A, Weiner OD, Lim WA, Voigt CA. Spatiotemporal control of cell signalling using a light-switchable protein interaction. Nature. 2009;461: 997–1001. doi:10.1038/nature08446

9. Strickland D, Lin Y, Wagner E, Hope CM, Zayner J, Antoniou C, et al. TULIPs: tunable, light-controlled interacting protein tags for cell biology. Nat Methods. 2012;9: 379–384. doi:10.1038/nmeth.1904

10. Guntas G, Hallett RA, Zimmerman SP, Williams T, Yumerefendi H, Bear JE, et al. Engineering an improved light-induced dimer (iLID) for controlling the localization and activity of signaling proteins. Proc Natl Acad Sci USA. 2015;112: 112–117. doi:10.1073/pnas.1417910112

11. O’Shaughnessy EC, Stone OJ, LaFosse PK, Azoitei ML, Tsygankov D, Heddleston JM, et al. Software for lattice light-sheet imaging of FRET biosensors, illustrated with a new Rap1 biosensor. J Cell Biol. 2019;218: 3153–3160. doi:10.1083/jcb.201903019

12. Liu B, Hobson CM, Pimenta FM, Nelsen E, Hsiao J, O’Brien T, et al. VIEW-MOD: a versatile illumination engine with a modular optical design for fluorescence microscopy. Opt Express. 2019;27: 19950–19972. doi:10.1364/OE.27.019950

13. Hodgson L, Nalbant P, Shen F, Hahn K. Imaging and photobleach correction of Mero-CBD, sensor of endogenous Cdc42 activation. Meth Enzymol. 2006;406: 140–156. doi:10.1016/S0076-6879(06)06012-5

14. Tsygankov D, Bilancia CG, Vitriol EA, Hahn KM, Peifer M, Elston TC. CellGeo: a computational platform for the analysis of shape changes in cells with complex geometries. J Cell Biol. 2014;204: 443–460. doi:10.1083/jcb.201306067

15. Caloca MJ, Wang H, Kazanietz MG. Characterization of the Rac-GAP (Rac-GTPase-activating protein) activity of beta2-chimaerin, a “non-protein kinase C” phorbol ester receptor. Biochem J. 2003;375: 313–321. doi:10.1042/BJ20030727

16. Iwig JS, Vercoulen Y, Das R, Barros T, Limnander A, Che Y, et al. Structural analysis of autoinhibition in the Ras-specific exchange factor RasGRP1. Elife. 2013;2: e00813. doi:10.7554/eLife.00813

17. Rubinfeld B, Munemitsu S, Clark R, Conroy L, Watt K, Crosier WJ, et al. Molecular cloning of a GTPase activating protein specific for the Krev-1 protein p21rap1. Cell. 1991;65: 1033–1042. doi:10.1016/0092-8674(91)90555-d

18. Michiels F, Habets GG, Stam JC, van der Kammen RA, Collard JG. A role for Rac in Tiam1-induced membrane ruffling and invasion. Nature. 1995;375: 338–340. doi:10.1038/375338a0

19. Kawasaki H, Springett GM, Toki S, Canales JJ, Harlan P, Blumenstiel JP, et al. A Rap guanine nucleotide exchange factor enriched highly in the basal ganglia. Proc Natl Acad Sci USA. 1998;95: 13278–13283. doi:10.1073/pnas.95.22.13278

20. Mao M, Wang L, Chang C-C, Rothenberg KE, Huang J, Wang Y, et al. Involvement of a Rac1-Dependent Macropinocytosis Pathway in Plasmid DNA Delivery by Electrotransfection. Mol Ther. 2017;25: 803–815. doi:10.1016/j.ymthe.2016.12.009

21. Harper SM, Neil LC, Gardner KH. Structural basis of a phototropin light switch. Science. 2003;301: 1541–1544. doi:10.1126/science.1086810

22. Crosson S, Moffat K. Photoexcited structure of a plant photoreceptor domain reveals a light-driven molecular switch. Plant Cell. 2002;14: 1067–1075. doi:10.1105/tpc.010475

23. Swartz TE, Corchnoy SB, Christie JM, Lewis JW, Szundi I, Briggs WR, et al. The photocycle of a flavin-binding domain of the blue light photoreceptor phototropin. J Biol Chem. 2001;276: 36493–36500. doi:10.1074/jbc.M103114200

24. Jaśkiewicz A, Pająk B, Orzechowski A. The many faces of rap1 gtpase. Int J Mol Sci. 2018;19. doi:10.3390/ijms19102848

25. Costa C, Barberis L, Ambrogio C, Manazza AD, Patrucco E, Azzolino O, et al. Negative feedback regulation of Rac in leukocytes from mice expressing a constitutively active phosphatidylinositol 3-kinase gamma. Proc Natl Acad Sci USA. 2007;104: 14354–14359. doi:10.1073/pnas.0703175104

26. de Beco S, Vaidžiulytė K, Manzi J, Dalier F, di Federico F, Cornilleau G, et al. Optogenetic dissection of Rac1 and Cdc42 gradient shaping. Nat Commun. 2018;9: 4816. doi:10.1038/s41467-018-07286-8

27. Wang H, Vilela M, Winkler A, Tarnawski M, Schlichting I, Yumerefendi H, et al. LOVTRAP: an optogenetic system for photoinduced protein dissociation. Nat Methods. 2016;13: 755– 758. doi:10.1038/nmeth.3926

