## Supplementary figures and images for "Optogenetic inhibition and activation of Rac and Rap1 using a modified iLID system"

### Supplemental Fig 1

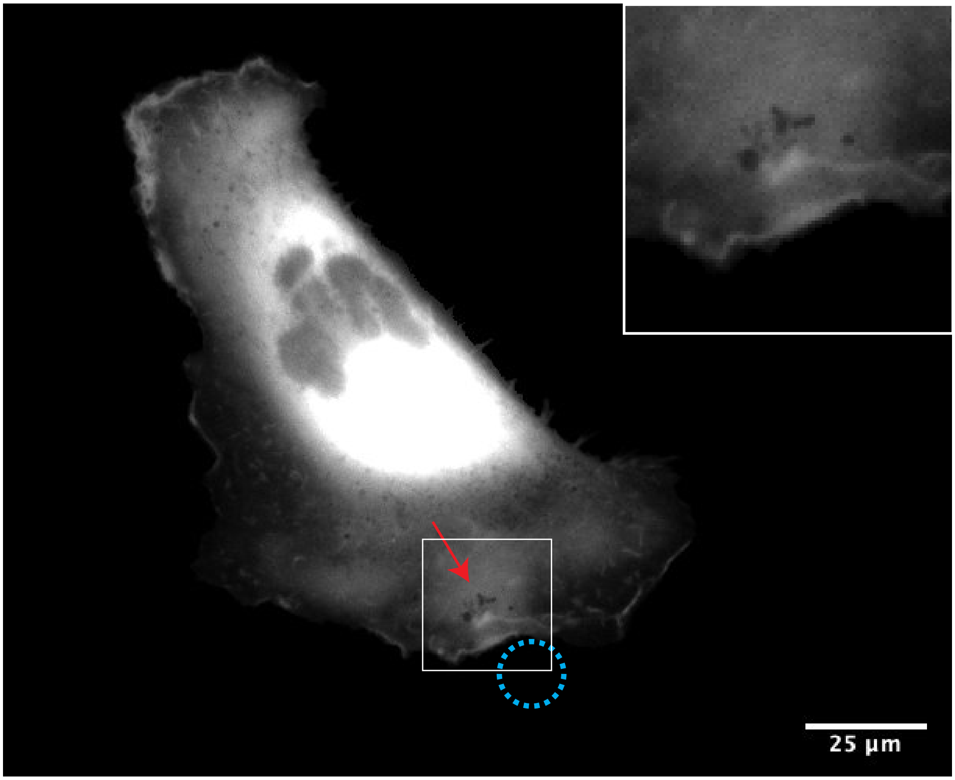


**Supplemental Figure 1.**
